# Absence of Replication fork associated factor CTF4 and F-box motif Encoding Gene SAF1 leads to reduction in Cell Size and Stress Tolerance Phenotype in *S. cerevisiae*

**DOI:** 10.1101/664185

**Authors:** Meenu Sharma, Samar Singh, V. Verma, Narendra K Bairwa

## Abstract

Chromosome transmission fidelity factor, Ctf4 in *S. cerevisiae* associates with replication fork and helps in the sister chromatid cohesion. At the replication fork, Ctf4 links DNA helicase with the DNA polymerase. The absence of Ctf4 invokes replication checkpoint in the cells. The Saf1 of *S*.*cerevisiae* interacts with Skp1 of SCF-E3 ligase though F box-motif and ubiquitinates the adenine deaminase Aah1 during phase transition due to nutrient stress. The genetic interaction between the CTF4 and SAF1 has not been studied. Here we report genetic interaction between CTF4 and SAF1 which impacts the growth fitness and response to stress. The single and double gene deletions of SAF1 and CTF4 were constructed in the BY4741 genetic background. The strains were tested for growth on rich media and media containing stress causing agents. The *saf1*Δ*ctf4*Δ cells with reduced cell size showed the fastest growth phenotype on YPD medium when compared with the *saf1*Δ, *ctf4*Δ, *and WT*. The *saf1*Δ*ctf4*Δ cells also showed the tolerance to MMS, NaCl, Glycerol, SDS, Calcofluor white, H_2_O_2_, DMSO, Benomyl, and Nocodazole when compared with the s*af1*Δ, *ctf4*Δ, and WT cells. However, *saf1*Δ*ctf4*Δ cells showed the sensitivity to HU when compared with WT and *saf1*Δ. Based on these observations we suggest that SAF1 and CTF4 interact genetically to regulate the cell size, growth and stress response.

## Introduction

*Saccharomyces cerevisiae*, utilized as bio-factory for production of biochemical and for understanding of basic biological processes such as DNA replication, chromosome segregation, autophagy, apoptosis etc. Yeast also used as biocontrol agent, for biofuel production, for green chemicals and enzymes synthesis. Stress tolerant yeast species or mutants are important requisite for bioprocessing industries (Raveendran *et al*. 2018). Bioprocessing industries requires yeast strains, which can tolerate osmotic, oxidative, thermal, starvation, acid and alkali stress, chemical inhibitors, heavy metal toxicity stress etc. The bioprocess industry obtained, stress tolerant yeast either from extreme environment or employs laboratory-engineering methods (Deparis *et al*. 2017). However, naturally occurring stress tolerant yeast are very rare.

A stress causes the loss of viability in yeast and reduces bioprocessing performance. The mechanisms of stress response in yeast mediated by variety of pathways which includes, HOG1 pathway (Brewster and Gustin 2014), protein kinase pathway (Papadakis and Workman 2015), and common stress signaling pathways (Folch-Mallol *et al*. 2004), which suggest that stress tolerance is polygenic trait. In yeast absence of a major gene or hub on a pathway, utilized to generate stress resistant strains. Lack of nutrients is the most common stress faced by yeast in wild, laboratory and industrial set-up. Therefore, strains, which are fitter in low nutrient, are important for industry. The ubiquitin proteasome system regulates the proliferation of *Saccharomyces cerevisiae* cells during stress caused by nutrients availability (Finley *et al*. 1987). The nutrient deprivation induces stress, which leads cells to enter into the quiescent phase (Finley *et al*. 1987). The SCF E3-ligase of ubiquitin proteasome system, recruits the substrate through the F-box-encoding gene for ubiquitination and subsequent degradation by 26S proteasome. During nutrient deprivation, adenine deaminase Aah1, of *S*.*cerevisiae*, which converts adenine to hypoxanthine, is degraded by proteasome. The F-box motif containing Saf1 recruits the Aah1 for ubiquitination (Escusa *et al*. 2006; Escusa *et al*. 2007).

The Replication fork associated factor Ctf4 constitutes the part of eukaryotic replisome and well conserved from yeast to humans. It acts as hub, which couples replisome factors through their Ctf4-interacting-peptide or CIP-box to the replication fork (Villa *et al*. 2016). The Ctf4 connects the DNA helicase and Pol alpha, absence of it invokes the replication checkpoint (Tanaka *et al*. 2009). Ctf4 mutant exhibits increased level of mitotic recombination at both inter-and intra-chromosomal loci and showed large budded cells with nucleus in the neck region (Kouprina *et al*. 1992). The mammalian homologue of CTF4 gene, And-1, have been shown to interact with the Mcm10 protein which associates with Mcm 2-7 helicase there by suggesting the role of CTF4 in the replication initiation. The antibody meditated disruption of the Mcm10- and And-1 interaction leads to defect in the loading of And-1 and DNA polymerase alpha to replication fork (Abe *et al*. 2018). Ctf4 homologue in fission yeast Mcl1 regulates S phase and deletion of it leads to cohesion defects (Williams and Mcintosh 2002). It interacts with F-box protein called Pof3, which belongs to SCF ubiquitin ligase complex. The mutant cells of pof3+ or mcl1+ showed accumulation of DNA damage and activation of DNA damage pathway (Mamnun *et al*. 2006). ctf4 mutant showed sensitivity to DNA damaging agents such as hydroxyurea (HU), phleomycin, camptothecin, and methyl methane sulfonate (MMS) similar to rad52 mutant suggesting a role of Ctf4 in recombination repair (Ogiwara *et al*. 2007). Genome wide genetic interaction studies reported CTF4 as hub which exhibits both positive and negative genetic interaction with large number of candidate genes (Collins *et al*. 2007; Costanzo *et al*. 2016; Kuzmin *et al*. 2018). However, SAF1 interacted genetically with CDC 10, CDC11, CDC12, HYP2 (Costanzo *et al*. 2016) negatively and showed positive interaction with CTF8 (Sharma et *al*. 2019, biorxiv archived data; doi 1101, www.biorxiv.org) only. Null mutant of SAF1 showed, synthetic growth defects with HSP82 (Zhao *et al*. 2005), POL2 (Dubarry *et al*. 2015), RTT109 (Fillingham *et al*. 2008) and RRM3 (Sharma *et al*. 2019, biorxiv archived data; doi 1101/636902, www.biorxiv.org).

Here we report the binary genetic interaction between F box motif encoding gene SAF1 and CTF4. Double deletion of SAF1 and CTF4 together leads to reduction in cell size, faster growth rate, and tolerance to wide range of stress causing agents except hydroxyurea in comparison to single gene mutant or WT.

## EXPERIMENTAL PROCEDURES

### Yeast strains and plasmids

The yeast strains and their genotype used in this study are mentioned in (**Table 1)**. The lists of plasmids used are mentioned **Table 2**. The ORF replacement was carried out as mentioned (Longtine *et al*. 1998).

**Table 1:**
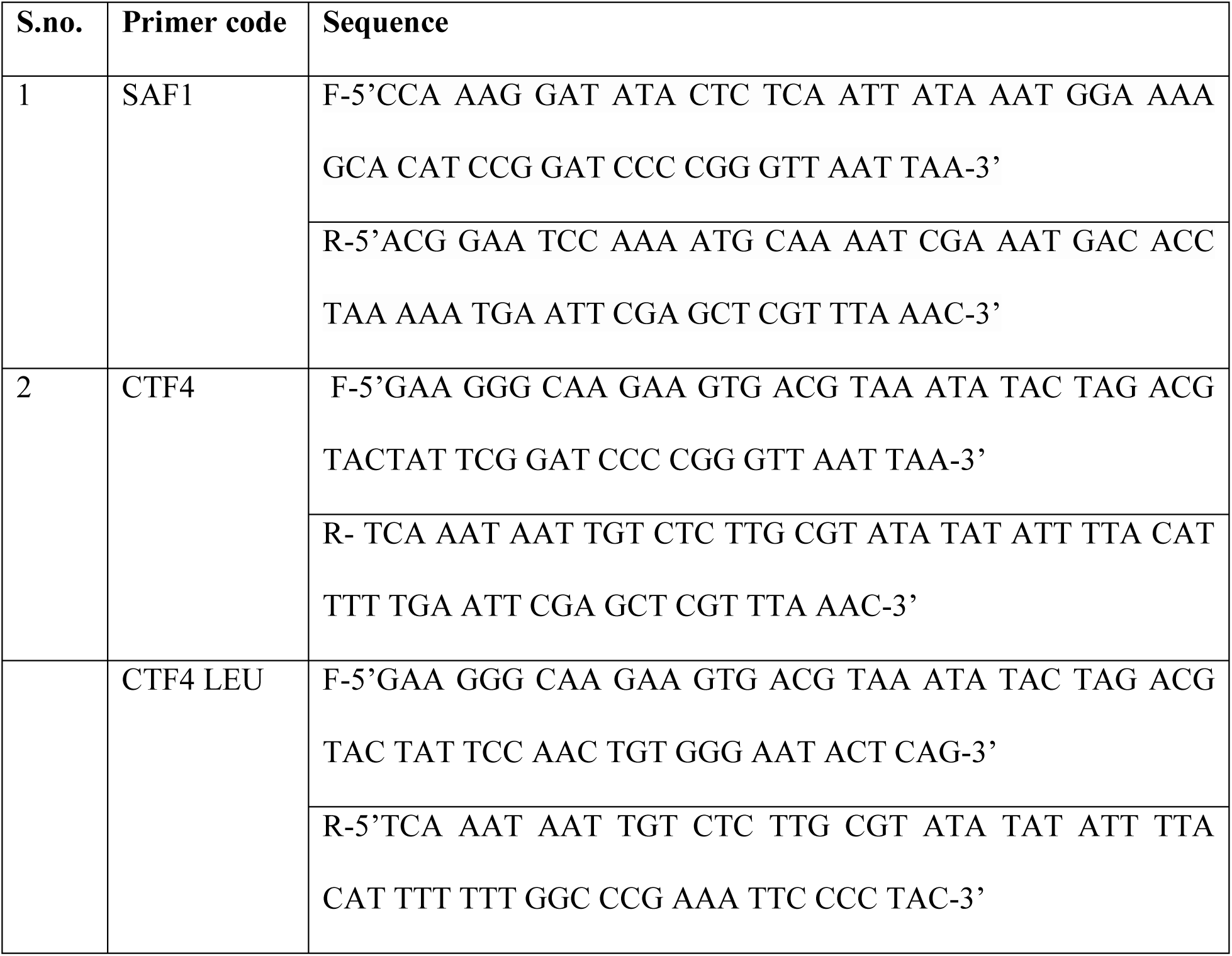
List of primers used for construction of deletion strains

**Table 2:**
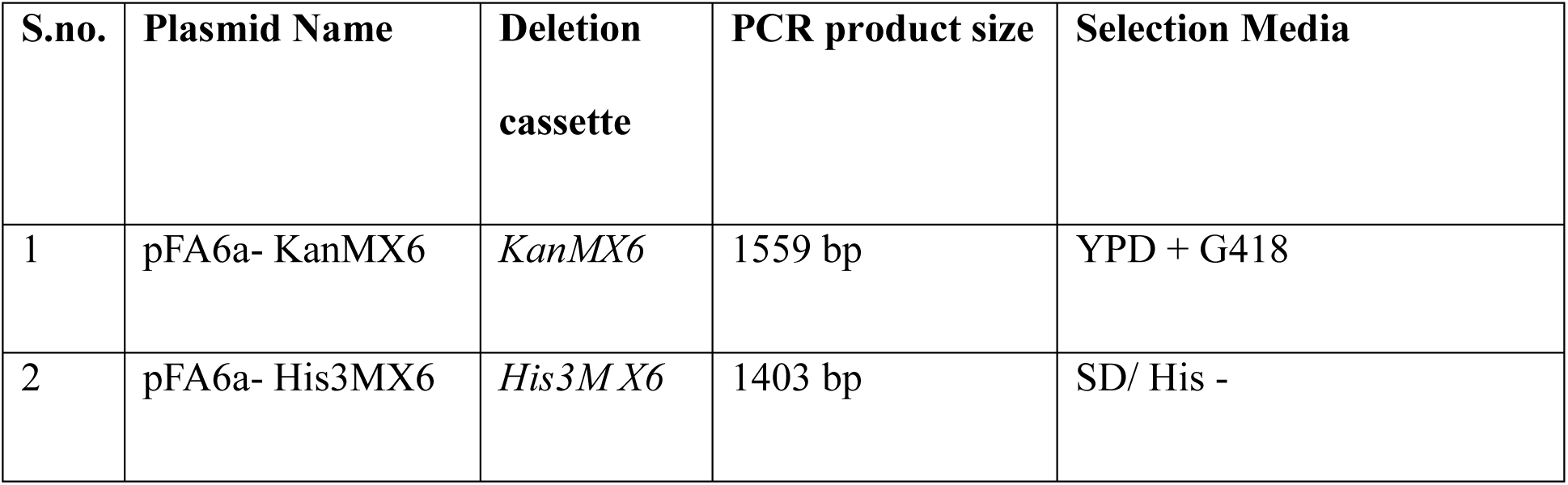
List of plasmids used for generating deletion cassette

**Table 3:** Yeast strains and their genotype used in the study

| S.no. | Strain | Genotype | Source |
| --- | --- | --- | --- |
| 1 | BY4741 | MATa his3Δ1 leu2Δ0 met15Δ0 ura3Δ0 | Dr. Deepak Sharma<br>IMTECH |
| 2 | MS1 | *saf1Δ::HIS3* | This study |
| 3 | MS2 | *ctf4Δ::KanMX* | This study |
| 4 | MS3 | *saf1Δ::HIS3, ctf4Δ::KanMX* | This study |
| 5 | JC2326 | MAT-ura3, cir0, ura3–167, leu::hisG, his32<br>Ty1his3AI-270, Ty1NEO-588, Ty1 (tyb::lacZ)-<br>146 | Prof. M. Joan<br>Curcio, USA |
| 6 | MJC1 | *saf1Δ::KanMX* | This study |
| 7 | MJC2 | *ctf4Δ::KanMX* | This study |
| 8 | MJC3 | *saf1Δ::KanMX, ctf4::LEU2* | This study |

### Growth Assay

Growth assessment of the WT and mutant strains on solid media was carried out by streaking on the YPD agar plates followed by incubation at 30°C for 2-3 days. For **g**rowth assessment of WT and mutants in the YPD broth, strains were grown for 14 hrs and optical density was measured every 2 hrs of interval at 600nm using TOSHVIN UV-800 SHIMADZU spectrophotometer. The OD values of three independent replicates of each culture were taken and average was plotted against time for growth curve.

### Phase Contrast Microscopy

To compare the morphology of WT and mutants, each **s**train was grown till the log phase in YPD medium at 30°C. The cultures were imaged by placing on slide under Leica DM3000 microscope at 100X magnification.

### Scanning Electron Microscopy (SEM)

For image acquisition of strains under scanning electron microscope, a single colony of each strain was inoculated in 10 ml YPD broth and grown for overnight in orbital shaker at 30°C at 180 rpm. Cells were suspended in 4% glutaraldehyde, prepared in 0.1M phosphate buffer, pH 7.2 and stored at 4°C for 1 hr. Cells were washed three times with 1 X PBS buffer and suspend in distilled water. Further cells were dehydrated through the ethanol series 30%, 50%, 70% and 95% wash. Finally, cells were suspended in 100% ethanol and dried at room temperature. Ethanol dried samples were mounted on to a SEM sample stub. Cells were sputtered with gold particles and viewed under the SEM, Model JSM 100 Jeol with image analyser.

### Calcofluor white staining and Fluorescence imaging

For staining of WT and mutant cells with Calcofluor white stain, method mentioned in (Pringle 1991; de Groot *et al*. 2001; Preechasuth *et al*. 2015) was adopted. Briefly, WT and each mutant strain were grown over night at 30°C and next day were re-inoculated in fresh YPD medium in 1:10 ratio. Cells were grown to log phase and collect by centrifugation. Collected cells were suspended in 100µl of solution containing Calcofluor white (50 µg/ml solution) fluorescent dye. Cells observed under 100X magnification using Leica DM3000 fluorescence microscope.

### Spot Assay

To assess growth fitness and cellular growth of WT and mutants in the presence of stress causing agents, spot assay was performed as mentioned in (Sharma et. al 2019, biorxiv archived data www.biorxiv.org). Briefly, BY4741 and its deletion derivatives strains *saf1*Δ, *ctf4*Δ, and *saf1*Δ*ctf4*Δ were grown in the 25 ml YPD (Yeast Extract 1% w/v, Peptone 2% w/v, dextrose 2% w/v) medium overnight at 30°C. The next day, overnight grown culture was diluted as 1:10 ratio in fresh YPD and grown until log phase (OD600 0.5-0.7). Equal OD value of cultures was adjusted and serially diluted. From each dilutions, an aliquot of 3µl was spotted onto agar plates containing YPD, YPD + stress causing agents such a Hydroxyurea (200mM), MMS (0.035%), SDS (0.0075%), Calcofluor (30µg/ml), 4% Glycerol, 1.4mM NaCl, 2mMH_2_O_2_, 8% DMSO. The plates were incubated at 30°C for 2-3 days and imaged. After recording of the cellular growth, cells from the first spotted lanes collected and observed under the100X magnification for image acquisition. The acquired images used for comparison of morphological features.

### Assay for Ty1 retro-mobility-

To measure the HIS3AI marked Ty1 retro-mobility, assay mentioned in (Scholes *et al*. 2001; Bairwa *et al*. 2011) and (Sharma et *al*. 2019, biorxiv archived data; doi 1101/636902, www.biorxiv.org) was performed. Briefly, WT (JC2326; reporter strain) and the deletion derivatives *saf1*Δ, *ctf4*Δ and *saf1*Δ*ctf4*Δ were inoculated into 10 ml YPD broth and grown overnight at 30°C. The overnight grown cultures were again inoculated in 5 ml YPD at 1:1000 ratios. The cultures were allowed to grow up to saturation point (144hrs) at 20°C. The saturated culture was serially diluted and plated on minimal media (SD/His^−^ plates) followed by incubation at 30°C for 3-7 days. The frequency of appearance of HIS^+^ colonies was measured for Ty1 retro-mobility.

### Statistical methods-

The significance of retro-mobility was determined using paired student t-test. P-value less than 0.05 indicated as significant.

## RESULTS

### Absence of both the genes SAF1 and CTF4 together leads to reduced cell size and faster growth phenotype

Both the genes SAF1 and CTF4 are non-essential in S. *cerevisiae*. Saf1 is involved in proteasome-dependent degradation of Aah1p during entry of cells into quiescence phase (Escusa *et al*. 2007). The null mutant of CTF4 showed slow growth rate in large-scale studies (Giaever *et al*. 2002) displayed the large cell size (Watanabe *et al*. 2009). We wished to determine the impact of deletion of both the gene together on growth fitness in rich medium. The single gene deletion of *saf1*Δ, *ctf4*Δ and double gene deletion, *saf1*Δ*ctf4*Δ were constructed in BY4741 genetic background. The strains were analysed for growth in YPD broth and on solid medium. We observed that *saf1*Δ and *ctf4*Δ showed slow growth in comparison to WT. However, *saf1*Δ *ctf4*Δ showed the fastest growth phenotype in comparison to WT, *saf1*Δ and *ctf4*Δ cells (**Figure 1 A, C**). The image comparison between WT, *saf1*Δ, *ctf4*Δ, *saf1*Δ*ctf4*Δ showed an enlargement of cell size in case of *ctf4*Δ (**Figure 1 B, Figure 2**) In contrast, *saf1*Δ *ctf4*Δ cells showed reduced cell size in comparison to WT, *saf1*Δ and *ctf4*Δ **(Figure 1 B, Figure 2)**. The observed phenotype of double mutant *saf1*Δ*ctf4*Δ in cell size reduction and fastest growth needs further exploration.

**Figure 1.**
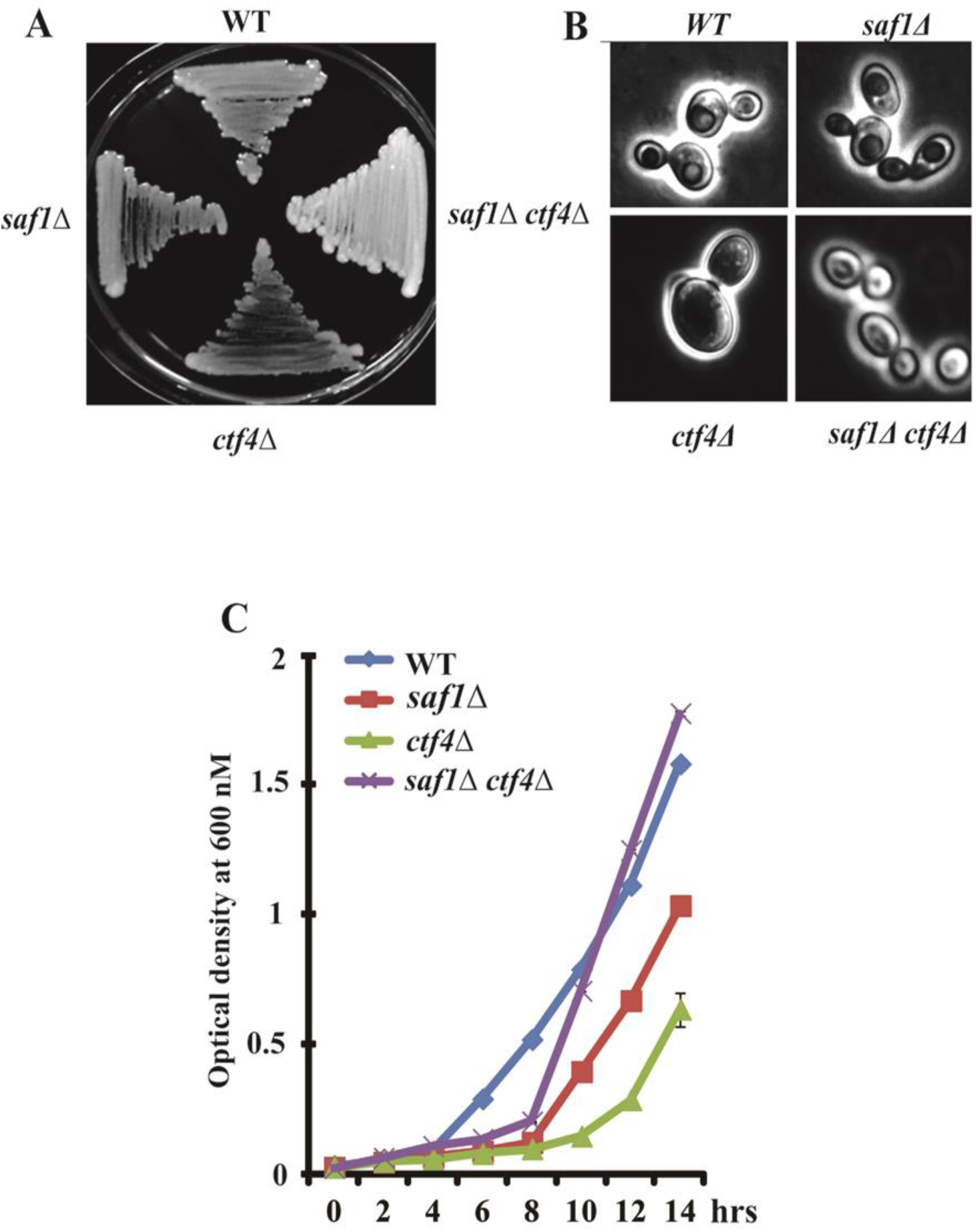
Comparative analysis of growth and morphology of WT, *saf1*Δ, *ctf4*Δ, *saf1*Δ*ctf4*Δ cells. **A.** Growth of streaked strains on YPD agar plates incubated for 2 days at 30°C and photographed. **B**. Phase contrast images of log phase cultures at 100X magnification using Leica DM3000. The *ctf4*Δ showed the enlargement of bud and mother cells whereas the *saf1*Δ*ctf4*Δ reduction in the cell size in comparison to WT, *saf1*Δ, *and ctf4*Δ cells. **C**. Growth kinetics of WT, *saf1*Δ, *ctf4*Δ, *saf1*Δ*ctf4*Δ cells, the double mutant cells showed the fastest growth in comparison WT, *saf1*Δ, *and ctf4*Δ. Cells were collected every 2 hour period and cellular growth was measured by optical density (OD) at 600 nm using TOSHVIN UV-1800 SHIMADZU. The data shown represent the average of three independent experiments. The error bars seen represent the standard deviation for each set of data.

**Figure 2.**
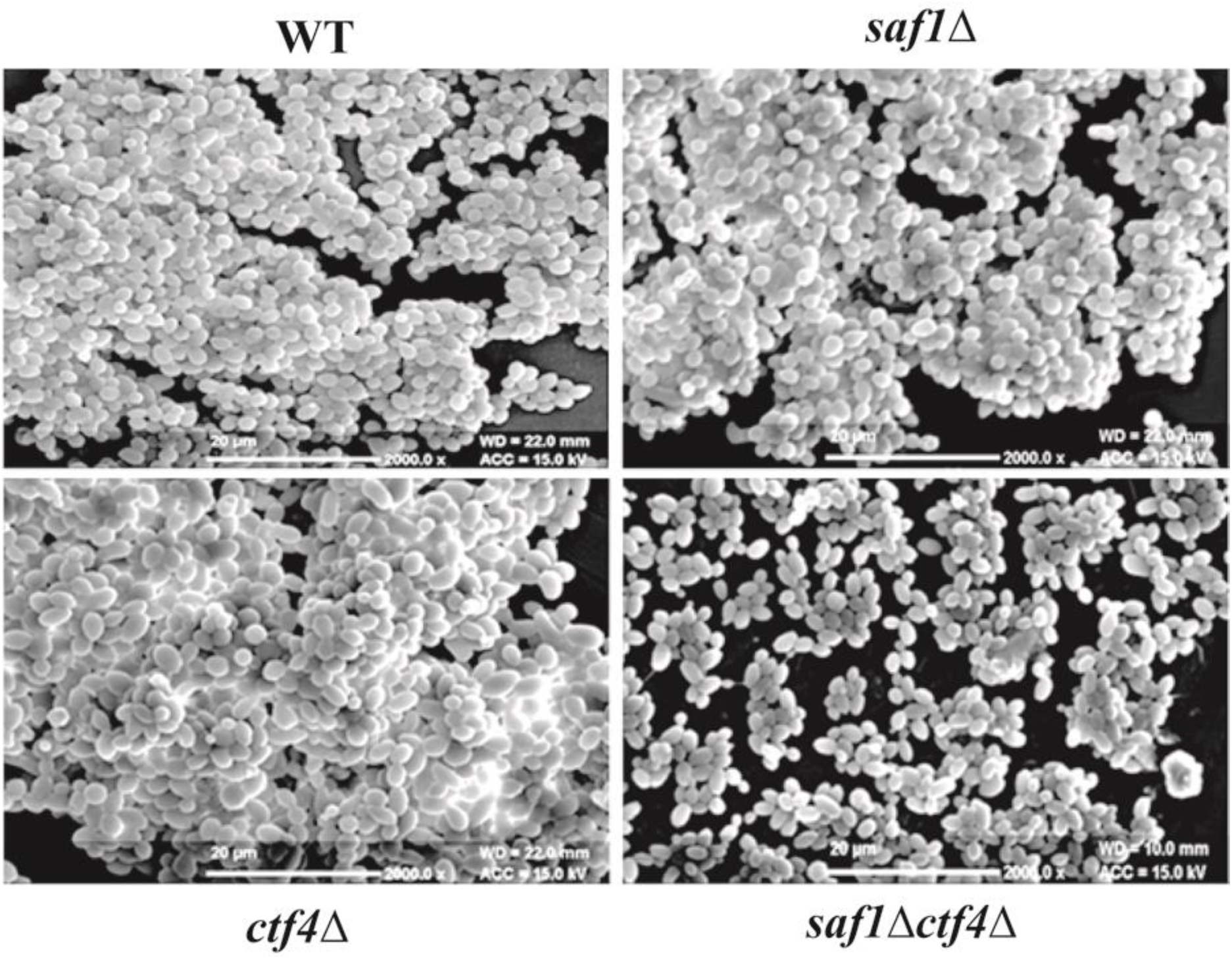
Comparative assessment of morphology and sizes of WT, *saf1*Δ, *ctf4*Δ, *saf1*Δ*ctf4*Δ strains using Scanning Electron Microscopy. The *ctf4*Δ cell showed the enlargement of bud and cell size whereas the *saf1*Δ*ctf4*Δ cells showed the reduction in size in comparison to WT, *saf1*Δ, and *ctf4*Δ cells.

### Loss of SAF1 and CTF4 together leads to MMS resistance and HU sensitivity

Methyl methane sulfonate (MMS) is DNA alkylating agent and elicit DNA damage in the cells after exposure. Hydroxyurea (HU) causes genotoxic stress by reducing the dNTP pool in the cell by inhibiting the activity of ribonucleotide reductase (RNR). We wished to study the cellular growth response of WT and *saf1*Δ, *ctf4*Δ, and *saf1*Δ*ctf4*Δ in the presence of 0.035% MMS and 200mM HU by semi-quantitative spot assay. The ctf4 mutant alone showed the sensitivity to hydroxyurea (HU), phleomycin, camptothecin, and methyl methane sulfonate (MMS) in earlier reported study (Ogiwara *et al*. 2007). In spot assay, *ctf4* showed extreme sensitivity to MMS and HU whereas *saf1*Δ*ctf4* cells showed resistance to MMS and sensitivity to HU in comparison to WT and *saf1*Δ (**Figure 3A, 3B)**. Further strains spotted on solid media having HU, showed the altered morphology depicting the defect in the mother daughter bud separation due to incomplete DNA replication (**Figure 3 C**). The observed cellular growth response to MMS and HU needs further investigation to understand the mechanism of DNA damage repair in the *saf1*Δ*ctf4*Δ mutant background.

**Figure 3.**
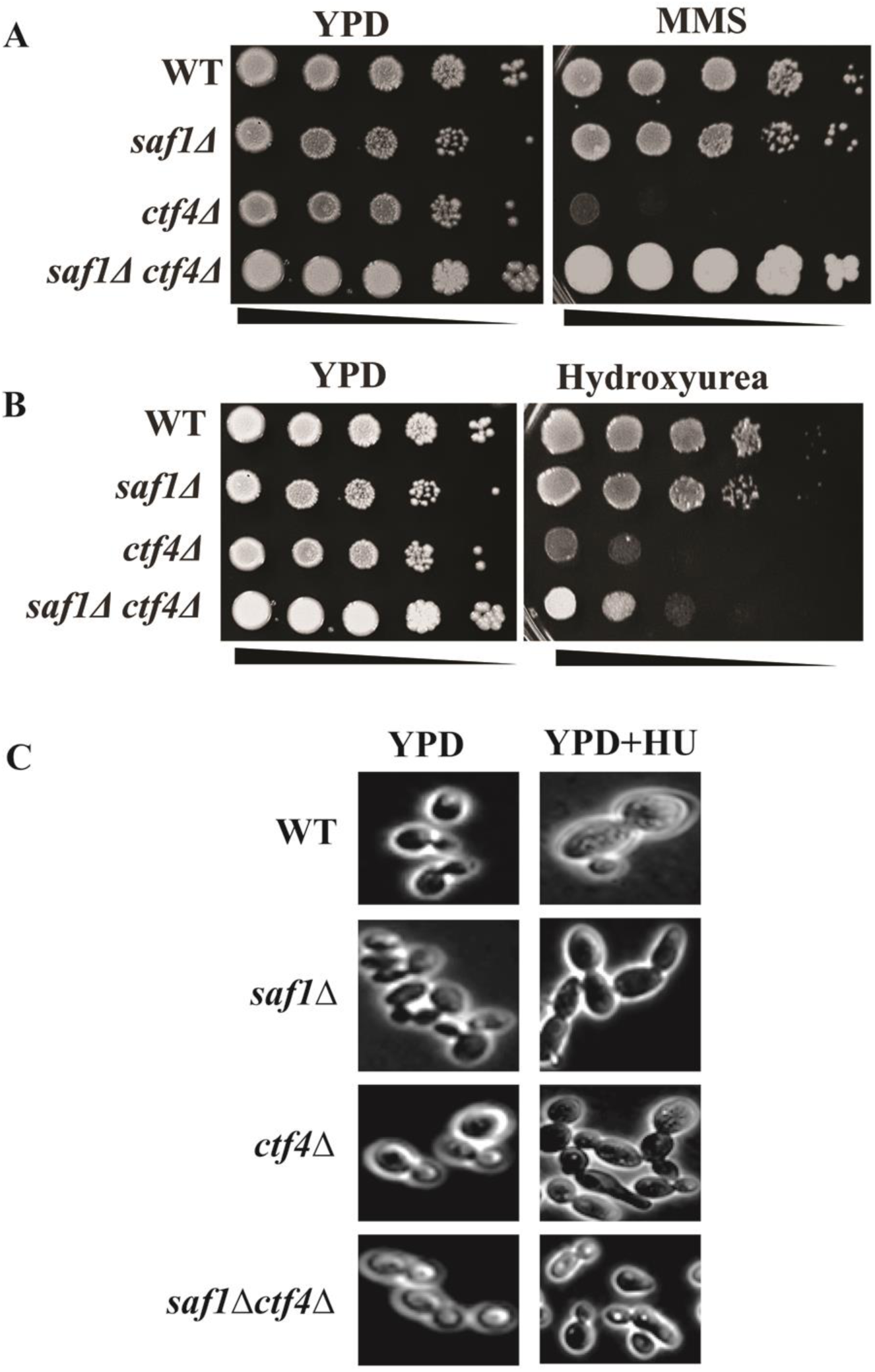
Comparative assessment of cellular growth response of WT, *saf1*Δ, *ctf4*Δ, *saf1*Δ*ctf4*Δ cells in presence of methyl methane sulfonate and hydroxyurea by spot analysis. Log phase culture equalized by O.D 600nm, serially diluted and spotted on YPD, YPD + HU (200mM) and YPD+MMS (0.035%) containing agar plates. A. The *saf1*Δ*ctf4*Δ showed resistance to MMS in comparison to WT, *saf1*Δ, *ctf4*Δ B. The *saf1*Δ*ctf4*Δ showed sensitivity to HU in comparison to WT, *saf1*Δ, *ctf4*Δ.

### Absence of SAF1 and CTF4 together contributes to Calcofluor white and SDS stress tolerance

Calcofluor white is a nonspecific flourochrome stain, which specifically binds to chitin and cellulose component of the cell wall (de Groot *et al*. 2001) and have been used as cell wall perturbing agents. Sodium dodecyl sulphate (SDS) acts as cell membrane disrupter. Both the agents have been used for screening of mutant as hypersensitive or resistance using spot assay. We wished to study the cellular growth response of WT and mutants (*saf1*Δ, *ctf4*Δ, and *saf1*Δ*ctf4*Δ) in the presence of 30µg/ml of Calcofluor white and 0.0075% SDS. Spot assay was carried out to observe the cellular growth response in presence of the stress-causing agents. We observed that WT, *saf1*Δ cells showed tolerance to Calcofluor white, whereas *saf1*Δ*ctf4*Δ showed slight sensitivity when compared with WT and *saf1*Δ *cells*. The c*tf4*Δ showed extreme sensitive phenotype the Calcofluor white (**Figure 4B**). Earlier studies reported *ctf4*Δ cells sensitive to Calcofluor white (Ando *et al*. 2007). The comparative image analysis of mutants showed the altered chitin distribution (**Figure 4A)**. The cellular growth response of mutants in presence of SDS showed double mutant *saf1*Δ*ctf4*Δ tolerant whereas *WT, saf1*Δ, *ctf4*Δ sensitive (**Figure 4C)**. It would be interesting to investigate the cell wall structure in saf1Δ*ctf4*Δ cells to understand the mechanism of Calcofluor white and SDS tolerance.

**Figure 4.**
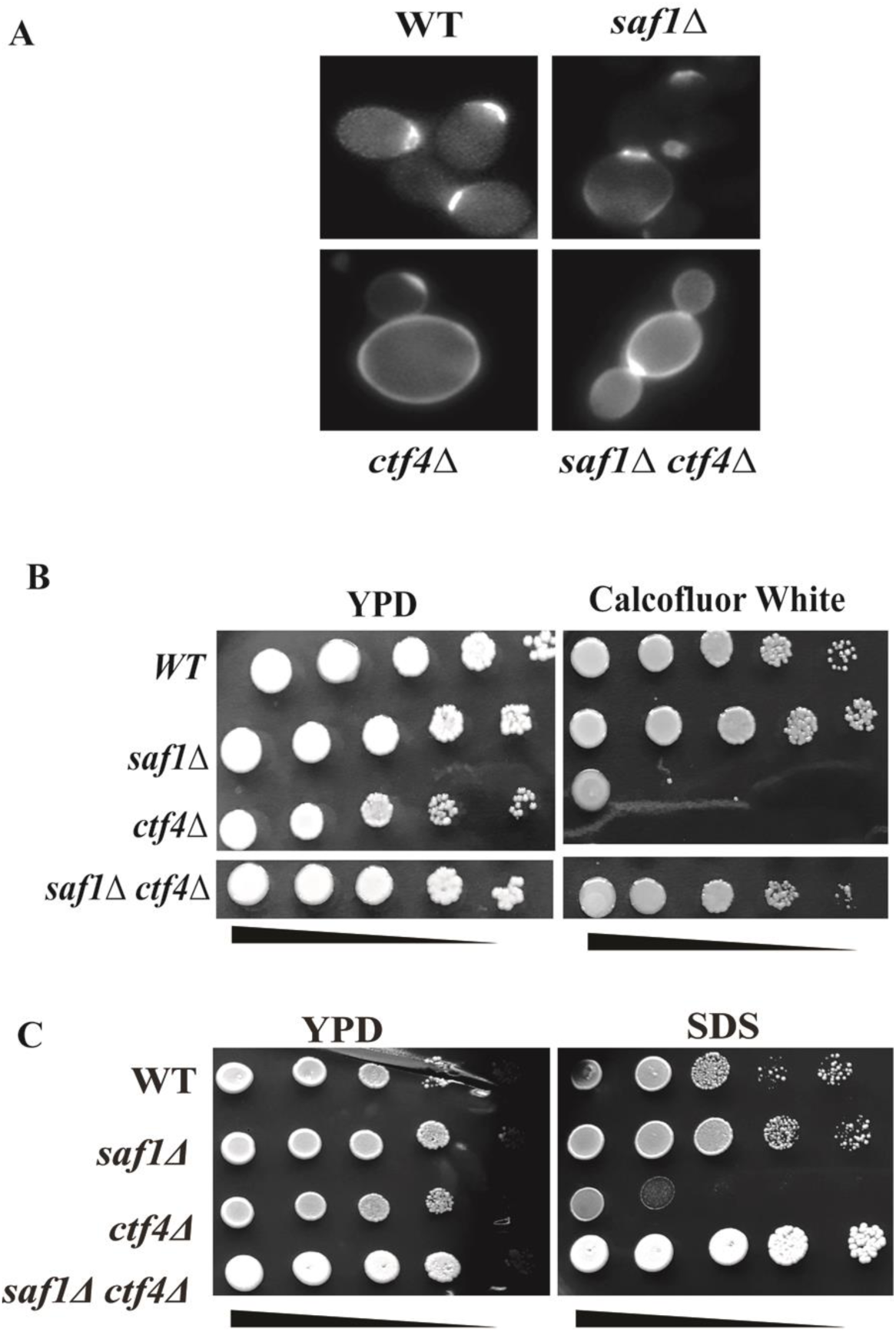
Comparative assessment of cellular growth response of WT, *saf1*Δ, *ctf4*Δ, *saf1*Δ*ctf4*Δ cells in presence of Calcofluor white and SDS by spot analysis. *C*alcofluor white stained cells imaged at 100X using Leica DM3000 fluorescence microscope. Log phase cultures equalized by O.D 600nm, serially diluted and spotted on YPD, YPD +Calcofluor white (30µg/ml) or SDS (0.0075%) containing agar plates. **A**. The *ctf4***Δ** showed cell enlargement and distributed chitin in comparison to WT and *saf1*Δ whereas *saf1*Δ*ctf4***Δ** showed reduction in size and distributed chitin in the cell wall. **B**. The *ctf4***Δ** showed the sensitivity to Calcofluor white whereas the *saf1***Δ***ctf4***Δ** showed resistance in comparison to *ctf4***Δ. C** The *ctf4***Δ** *cells* showed extreme sensitivity to SDS whereas the *saf1***Δ***ctf4***Δ** cells showed resistance in comparison to WT, *saf1***Δ** and *ctf4*Δ alone.

### Absence of SAF1 and CTF4 together leads to oxidative stress tolerance caused by DMSO and H_2_O_2_

Dimethyl sulfoxide (DMSO) is an amphiphilic compound, which contains the hydrophilic sulphoxide and hydrophobic methyl groups. The hydrophilic group defines the action of DMSO on the membrane and used as effective penetration enhancer and cryoprotectant (Sadowska-Bartosz *et al*. 2013). In *S. cerevisiae*, DMSO reported to inhibit the activity of methionine sulfoxide reductase A, thereby inhibiting the generation of methionine-S-sulfoxide (Kwak *et al*. 2010). The DMSO induces oxidative stress in yeast cells as reported in (Sadowska-Bartosz *et al*. 2013).The free radical generating compound’s exposure and aerobic metabolism both generate reactive oxygen species (ROS) in all organisms. The ROS are toxic functional group, which causes damage to cellular components including modification of DNA. An oxidative stress, characterized as when the antioxidant and cellular survival mechanisms both compromised following exposure to ROS. Hydrogen peroxide (H_2_O_2_) used as oxidative stress inducing agent. Here we wished to study the cellular growth response of WT and mutants (*saf1*Δ, *ctf4*Δ, and *saf1*Δ*ctf4*Δ) in the presence of 8% DMSO and 2mM hydrogen peroxide. We observed that WT, *saf1*Δ, *ctf4*Δ showed extreme sensitivity to DMSO. However saf1Δ*ctf4*Δ showed resistance to DMSO (**Figure 5A**). Incase exposure to H_2_O_2_ oxidative stress, *saf1*Δ*ctf4*Δ *cells* showed tolerance in comparison to WT, *saf1*Δ and *ctf4*Δ (**Figure 5B**). To understand the mechanism of tolerance to oxidative stress (DMSO and H_2_O_2)_ in *saf1*Δ*ctf4*Δ mutant needs further investigation.

**Figure 5.**
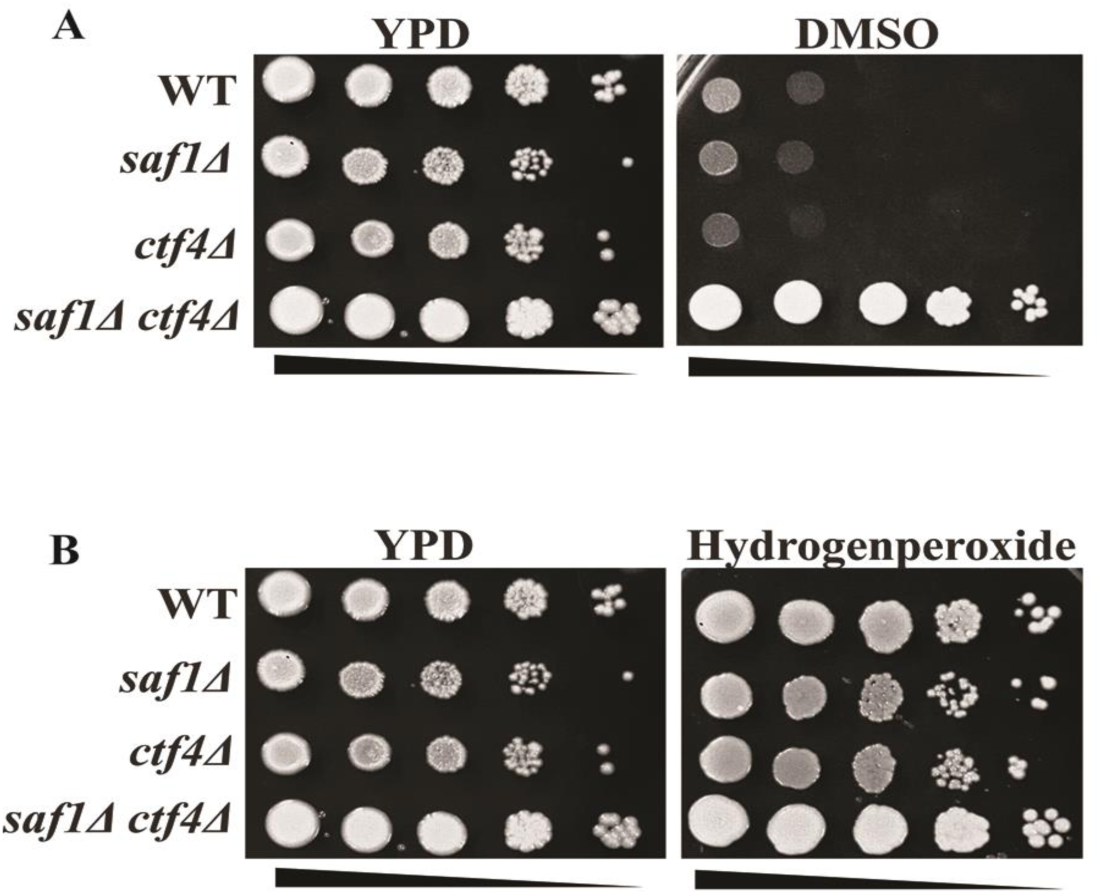
Comparative assessment of cellular growth response of WT, *saf1*Δ, *ctf4*Δ, *saf1*Δ*ctf4*Δ cells in presence of oxidative stress agents, DMSO and H_2_O_2_ by spot analysis. Log phase cultures equalized by O.D 600nm, serially diluted and spotted on YPD, YPD + DMSO (8%) or H_2_O_2_ (2mM) containing agar plates. **A**. The *saf1*Δ*ctf4*Δ cell showed the tolerance to DMSO whereas *WT, saf1***Δ** and *ctf4*Δ showed no growth. **B** The *saf1*Δ*ctf4*Δ cell showed the extreme tolerance to H_2_O_2_ in comparison to *WT, saf1***Δ** and *ctf4*Δ.

### Absence of SAF1 and CTF4 together leads to Osmotic stress tolerance caused by Glycerol and NaCl

Glycerol is an important constituent of yeast cells. It serves as carbon source, osmolyte and function as metabolite as its synthesis leads to regulation of cellular redox balance (Duskova *et al*. 2015). High concentrations of external glycerol allow cells to activate transient induction of the expression of stress protective genes, which leads to accumulation of intracellular glycerol. The tolerance of high concentration of glycerol by yeast is important for biotechnological applications. *S. cerevisiae* has been a great model for understanding mechanism of salt stress. When yeast cells exposed to saline stress they face both osmotic stress and cation toxicity. We investigated the cellular growth response of WT and mutants (*saf1*Δ, *ctf4*Δ, and *saf1*Δ*ctf4*Δ) in presence of 4% glycerol and 1.4M NaCl. We found that WT, *saf1*Δ, *ctf4*Δ showed slight sensitivity. However *saf1*Δ*ctf4*Δ showed resistance to 4% glycerol (**Figure 6A**). In case of salt stress, WT, *saf1*Δ, *ctf4*Δ failed to *g*row in presence of 1.4 M NaCl however *saf1*Δ*ctf4*Δ showed robust growth (**Figure 6B**). The mechanism of salt stress tolerance due to SAF1, CTF4 ablation needs further investigation.

**Figure 6.**
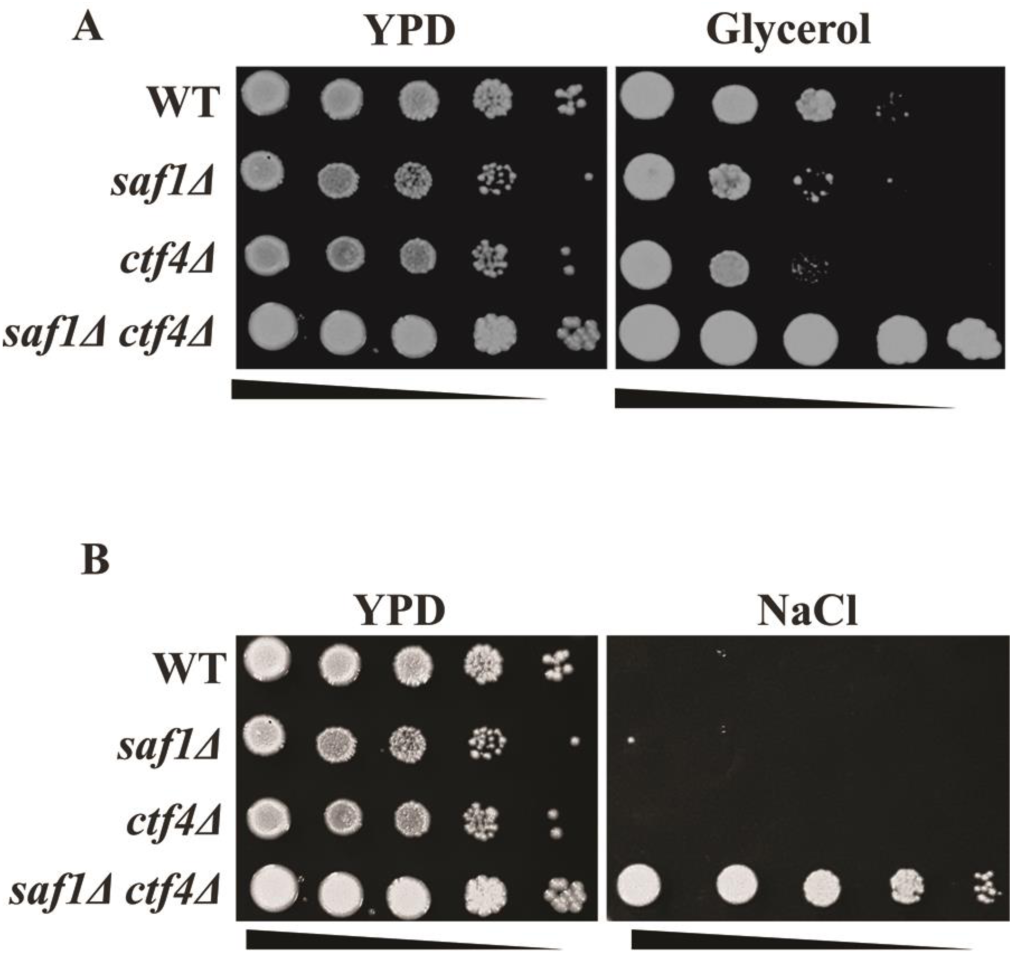
Comparative assessment of cellular growth response of WT, *saf1*Δ, *ctf4*Δ, *saf1*Δ*ctf4*Δ cells in presence of osmotic stress agents, glycerol and NaCl by spot analysis. Log phase cultures equalized by O.D 600nm, serially diluted and spotted on YPD, YP+ Glycerol (4%) or YPD + NaCl (1.4M) containing agar plates. **A**. The *saf1*Δ*ctf4*Δ cell showed the robust growth in presence of glycerol whereas *WT, saf1***Δ** and *ctf4*Δ showed reduced growth on glycerol containing plates. **B** The *saf1*Δ*ctf4*Δ cell showed growth in presence of NaCl in comparison to *WT, saf1***Δ** and *ctf4*Δ that showed no growth.

### Absence of SAF1 and CTF4 together leads to Benomyl and Nocodazole resistance

Microtubule depolymerizing agent, benomyl, affects movement of chromosomes and cell division. It binds to β-tubulin, coded by TUB2 gene and interferes in large number of cellular processes, which involves the participation of microtubules. The disruption in polymerization of microtubules affects the process of exit from mitosis and cause apoptosis (Thomas *et al*. 1985; Jordan and Wilson 2004). Nocodazole interacts with the free tubulin, which affect the cytoskeleton formation and nuclear division. We investigated the cellular growth response of WT and mutants (*saf1*Δ, *ctf4*Δ, and *saf1*Δ*ctf4*Δ) in presence of 100µg/ml benomyl and 50µg/ml Nocodazole. We observed that WT, *saf1*Δ, ctf4Δ showed sensitivity to benomyl and Nocodazole. In contrast, *saf1*Δ*ctf4*Δ showed the resistance (**Figure 7A and 8A**) to both the drugs. The image analysis of the cells exposed to the both the drugs showed characteristic elongated buds, in WT and single gene mutants (WT, *saf1*Δ, *ctf4*Δ) however, the double mutant (*saf1*Δ*ctf4*Δ) cells failed to show the elongated bud (**Figure 7B and 8B**).

**Figure 7.**
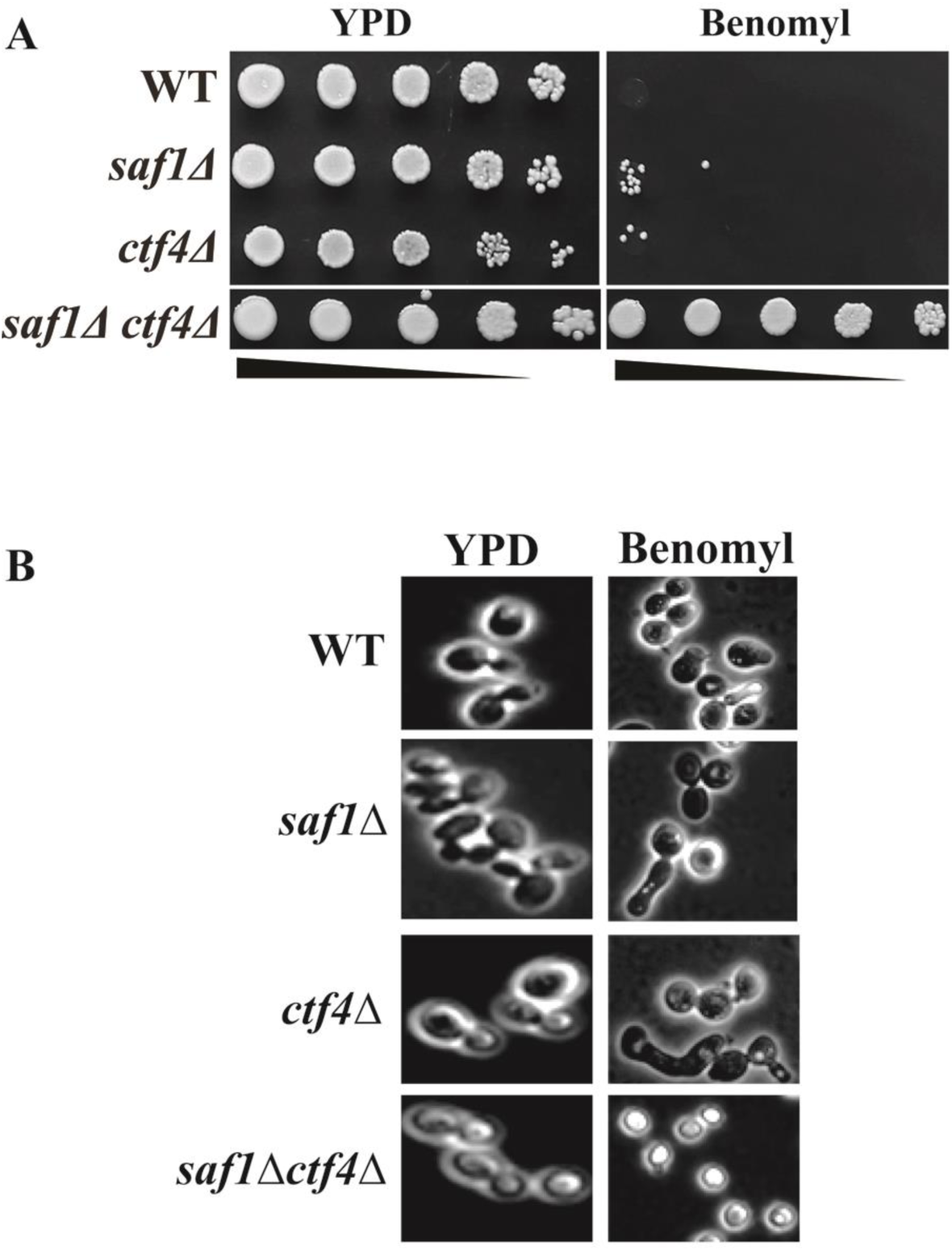
Comparative assessment of cellular growth response of WT, *saf1*Δ, *ctf4*Δ, *saf1*Δ*ctf4*Δ cells in presence of Microtubule depolymerizing drug Benomyl by spot analysis. Log phase cultures equalized by O.D 600nm, serially diluted and spotted on YPD, YPD+ Benomyl (100µg/ml) containing agar plates. **A**. The *saf1*Δ*ctf4*Δ cell showed growth and resistance to benomyl whereas *WT, saf1***Δ** and *ctf4*Δ showed extreme sensitivity to benomyl presence.**B**. Comparative phase contrast images of WT, *saf1*Δ, *ctf4*Δ, *saf1*Δ*ctf4*Δ cells grown in presence of microtubule depolymerizing drug benomyl. The WT, *saf1*Δ, *ctf4*Δ cells showed the characteristic elongated tube formation when treated with benomyl whereas the saf1Δ*ctf4*Δ *cells* showed no elongated tube formation

**Figure 8.**
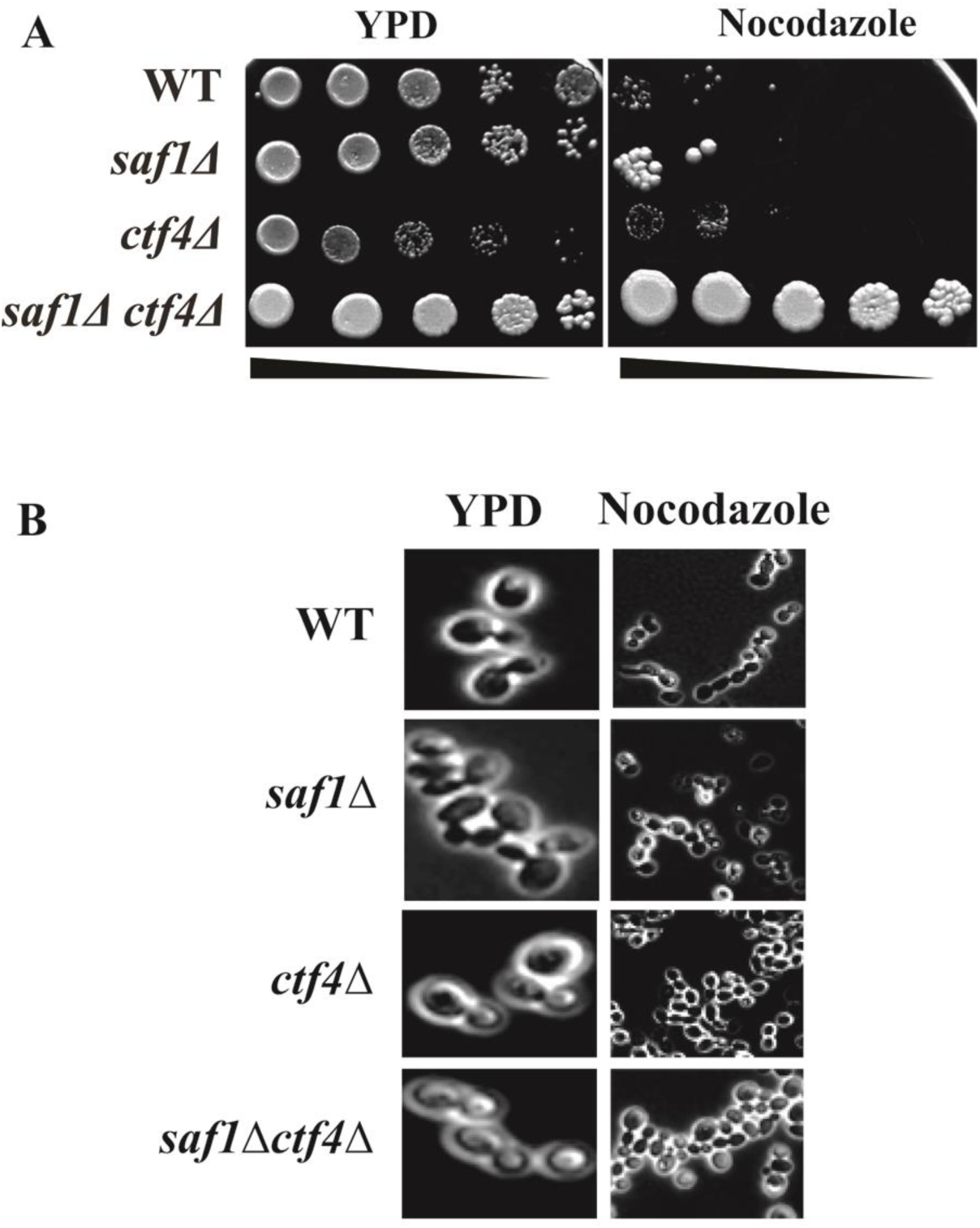
Comparative assessment of cellular growth response of WT, *saf1*Δ, *ctf4*Δ, *saf1*Δ*ctf4*Δ cells in presence of Nocodazole by spot analysis. Log phase cultures equalized by O.D 600nm, serially diluted and spotted on YPD, YPD+ Nocodazole (50µg/ml) containing agar plates. **A**. The *saf1*Δ*ctf4*Δ cell showed growth and resistance to Nocodazole whereas WT, *saf1***Δ** and *ctf4*Δ showed extreme sensitivity to Nocodazole. **B**. Comparative phase contrast images of WT, *saf1*Δ, *ctf4*Δ, *saf1*Δ*ctf4*Δ cells grown in presence of Nocodazole. The WT, *saf1*Δ, *ctf4*Δ cells showed the characteristic chained cells indicating of G2-M arrest whereas the saf1Δ*ctf4*Δ cells showed budding pattern.

### Deletion of both SAF1 and CTF4 induced high frequency of Ty1 retro–transposition

Ty1 retro-mobility in S. *cerevisiae* is induced by *r*eplication stress and DNA damage. The absence of the ctf4 leads to replication stress and cohesion defects. The replication stress in S-phase leads to checkpoint induction and increased Ty1 retro-transposition as reported earlier (Curcio *et al*. 2007; Bairwa *et al*. 2011).The absence of SAF1 promotes the Ty1 retro-transposition (Sharma et *al*. 2019, biorxiv archived data; doi 1101/636902, www.biorxiv.org). Here we observed that *saf1*Δ (upto 8-10 fold) and *ctf4*Δ, *(nearly* 75 fold) showed increase in the Ty1 retro-transposition in comparison to WT (**Figure 9A, B**), however *saf1*Δ*ctf4*Δ showed, 150 fold increase in the Ty1 retro-transposition **(Figure 9A, B**). These observations indicate that SAF1 and CTF4 requires for suppression of Ty1 retro-transposition.

**Figure 9.**
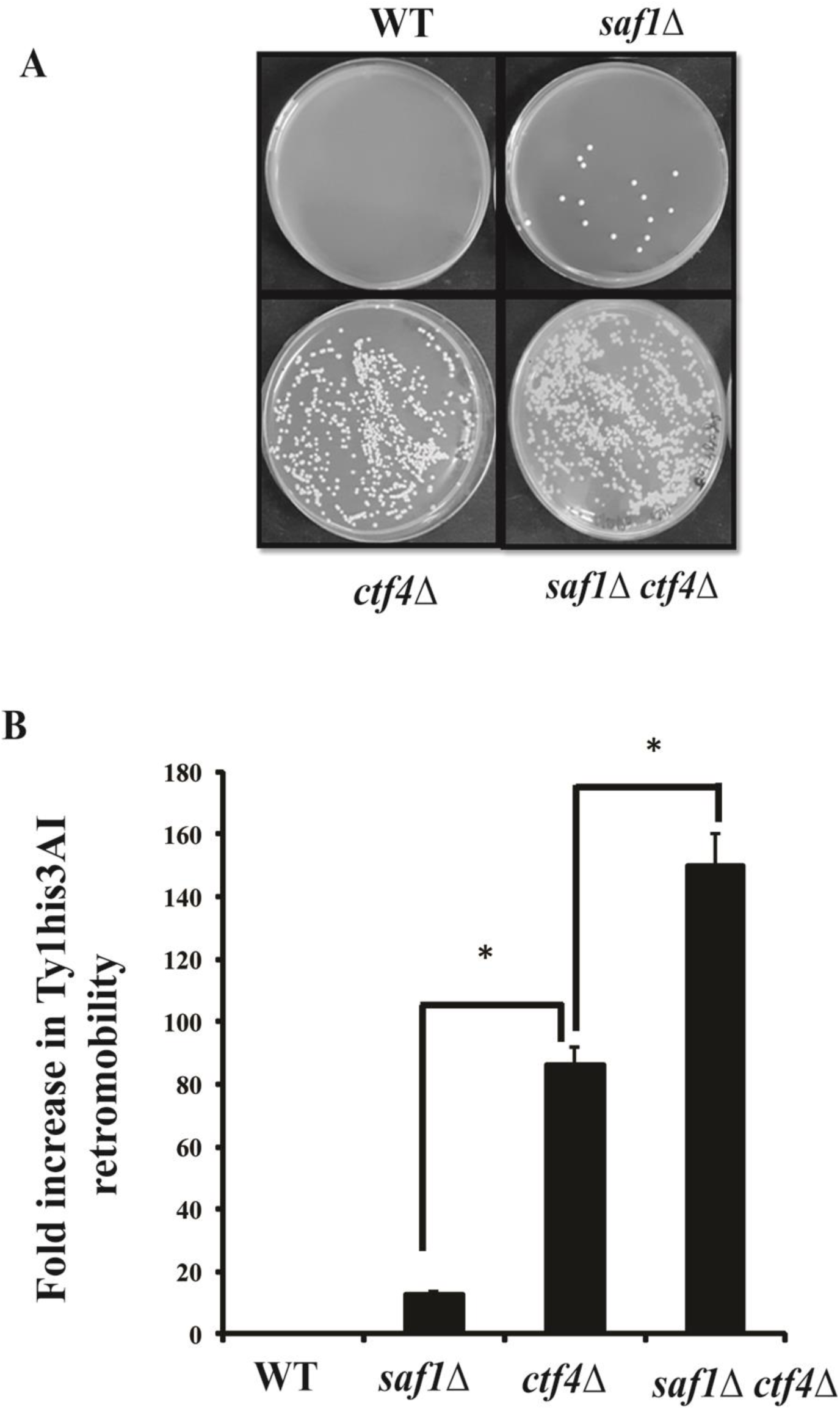
Comparative assessment of HIS3AI marked Ty1 transposition frequency in WT, *saf1*Δ, *ctf4*Δ, *and saf1*Δ*ctf4 strains*. **A** Images of plates showing the Ty1 transposition induced colonies on SD plate lacking His media. **B**. Bar diagram showing the frequency of Ty1his3AI transposition in each strain. The data shown represent the average of three independent experiments. The significance of transposition was determined by using two tailed t-test. P-value (p) less than 0.05 indicate significant difference and the symbol * represent to p<0.05.

## Discussion

Here, we have investigated genetic interaction between the SAF1 and replication fork associated factor CTF4. The binary genetic interactions studies helps in understanding the roles of individual genes in the compensatory pathways, which regulates the biological processes. This understanding helps in building of gene networks for system biology applications. In this work, we discovered positive genetic interaction between SAF1 and CTF4 genes, which regulates the growth fitness, and cell size in the *S. cerevisiae*. We also show that cells lacking both the genes, tolerates wide range of stressor such as DMSO, H2O2, SDS, Calcofluor white, Glycerol, NaCl, MMS, Nocodazole, and Benomyl, the only stress causing agent which cell did not tolerate well was hydroxyurea. Our data revealed that both the genes work in compensatory pathway.

The SAF1 showed synthetic growth defect with HSP82, HSC82 (McClellan *et al*. 2007), RTT101 (Fillingham *et al*. 2008), POL2 (Dubarry *et al*. 2015), IZH2 (Mattiazzi Usaj *et al*. 2015) and with DNA helicase RRM3 (Sharma et *al*. 2019, biorxiv archived data; doi 1101/636902; www.biorxiv.org). The synthetic rescue phenotype of SAF1 has been observed with the NPL3, whose product helps in co-transcriptional recruitment of the splicing machinery (Moehle *et al*. 2012), ESS1, which codes for prolyl isomerase regulates the nuclear localization of Swi6 and Whi5 (Atencio *et al*. 2014). The CTF8 gene, product constitutes the part of RFC-Ctf18 complex and helps in loading of PCNA on to the chromosome, showed positive interaction with SAF1 (Sharma et *al*. 2019, biorxiv archived data; doi 1101 www.biorxiv.org). Here we discovered that SAF1 deletion rescued the growth defects of cells lacking the CTF4. Other genes ASF1, MMS1, MMS22, RRM3, RTT101, RTT109 (Luciano *et al*. 2015) SET2 (Biswas *et al*. 2008), POB3 (Schlesinger and Formosa 2000), HST4, HST3 (Celic *et al*. 2008; Che *et al*. 2015), FOB1 (Budd *et al*. 2005; Shyian *et al*. 2016; Sasaki and Kobayashi 2017) and DIA2 (Pan *et al*. 2006) also have been shown to restore the normal growth of CTF4 null mutant. Ctf4 protein acts as hub, linking DNA helicase and DNA polymerase with other many factors, which are involved in other metabolic processes. The cell lacking CTF4 alone showed extreme to moderate sensitive phenotype when grown in presence of stress causing agents *i*.*e*. Hydroxyurea (Parsons *et al*. 2004), MMS (McKinney *et al*. 2013), Cycloheximide (Dudley *et al*. 2005), Calcofluor white (Ando *et al*. 2007; Kapitzky *et al*. 2010), Benomyl (Daniel *et al*. 2006) and NaCl (Michaillat and Mayer 2013). In our study, null mutant of CTF4 also showed the moderate to extreme sensitive phenotype in presence of the HU, MMS, and Calcofluor white, SDS, H_2_O_2_, NaCl, Benomyl and Nocodazole suggesting its role in linking of various biological pathways involved in stress response to replication fork components. The null mutant SAF1 showed resistance to histone deacetylase inhibitor CG-1521 drug (Gaupel *et al*. 2014). In our study null mutant of SAF1 showed the cellular growth in presence of, H_2_O_2,_ Calcofluor white, SDS, MMS, and HU whereas sensitivity to the DMSO and NaCl. This suggests that Saf1 function differently in response to variety of stress. Further, increased frequency of Ty1 retro-transposition in double mutant of SAF1, CTF4 suggest both are required for transcriptional dormancy of the Ty1 element. The tolerance of *saf1*Δ*ctf4*Δ cells to stress causing agents except hydroxyurea is the novel outcome of the study. Further it would be interesting to investigate into the contribution of hCTF4/AND1 and HERC2 for metastatic potential and drug resistance in the cancer cell lines or tumours. We also suggest that an investigation may be undertaken to understand the mechanism of drug resistance and contribution of SAF1 and CTF4 homologues in pathogenic fungal species. The combined observation, on cell size reduction, fastest growth, salt and other stress tolerance by *saf1*Δ*ctf4*Δ cells need to be extended to other model organism as both the genes are functionally well conserved from yeast to humans. This may yield novel phenotypes and biotechnological applications.

## Acknowledgment

We thank to Prof. M. Joan Curio, Dr. Deepak Sharma, IMTECH for strains and plasmids. We also thank Dr. Jitendra Thakur, NIPGR for his Lab support and Panjab University for SEM facility.

## Author’s contributions

NKB conceived and directed the study and wrote the paper with MS. MS performed the experiments and analysed data with NKB. SS and VV provided the SEM and bioinformatics facility and analysed the data. All the authors reviewed the results and approved the final version of manuscript.

